# From the Wild to the City: How Domestication and Urbanization Reshape Animal Gut Microbiome

**DOI:** 10.1101/2023.11.20.567780

**Authors:** Angsuman Das, Bhabana Das, Jyotishmita Das

## Abstract

This review explores the profound effects of domestication and urbanization on the gut microbiota of animals. It delves into the complex interplay between these two processes and their transformative impact on the microorganisms residing in the gastrointestinal tracts of a wide range of species. Domestication, the centuries-old practice of taming and breeding animals for human use, has led to significant shifts in the gut microbiomes of domesticated animals. This shift is a result of altered diets, living conditions, and reduced exposure to natural environments. The paper examines the consequences of these changes on animal health, behavior, and their adaptation to domestic life. Conversely, urbanization, characterized by the rapid expansion of cities and human habitats, has driven wild animals to adapt to urban environments. This review investigates how the urban landscape, pollution, and dietary changes reshape the gut microbiomes of urban wildlife. It explores the potential implications of these alterations on the animals’ resilience to urban stressors and disease. Drawing parallels between domestication and urbanization, the paper reveals intriguing similarities and differences in gut microbiome transformations across various species. It also assesses the broader implications of these shifts on ecological dynamics, zoonotic disease transmission, and the potential for microbial interactions between domesticated animals, urban wildlife, and humans. Ultimately, this review consolidates current knowledge on the topic, shedding light on the shared mechanisms and unique adaptations that drive microbial changes in animals undergoing domestication and those adapting to urban environments. It concludes with a discussion of the implications for animal conservation, animal-human interactions, and the One Health perspective, emphasizing the importance of understanding these intricate icrobial relationships in our ever-changing world. By enhancing our comprehension of these complex dynamics, this paper contributes to the growing body of knowledge that informs our coexistence with the animals we share our lives and cities with, highlighting the critical role of gut microbiota in these processes.

## I. Introduction

The gut microbiome plays a fundamental role in the health and evolution of both humans and animals. It is involved in digestion, immunity, and even behavior. The composition of the gut microbiome is influenced by various factors such as age, race, gender, geographical distribution, and genetic features (Park et al., 2023) (D’Amico et al., 2023). Dysbiosis of the gut microbiota has been associated with the development and progression of age-related disorders in the elderly population (Davenport et al., 2017). Mounting evidence suggests that the gut microbiome is implicated in the development of a wide range of diseases, including gastrointestinal diseases, metabolic diseases, neurological disorders, and cancers (Thriene & Michels, 2023). The plasticity of the gut microbiota is highest at birth and decreases with advancing age, which has implications for the impact of the exposome on the microbiota and health throughout the life course (Bastón-Paz et al., 2022)[5]. Alterations in gut microbiota parameters have been associated with cardiovascular comorbidities, obesity, and type 2 diabetes, highlighting the potential for modifying the microbiome to achieve clinical benefits.

The transition from hunter-gatherer societies to agricultural settlements and then to industrial urbanization was a significant turning point in human history. During the Mesolithic period, societies relied on hunting and gathering for sustenance (Szabo & Szabo, 2016). However, the development of agriculture in the Neolithic period brought about major changes in economic base, material culture, settlement patterns, and population levels (Dabrowski & Haynor, 2017). This transition was accompanied by the domestication of animals, which played a crucial role in agricultural practices (Innes & Blackford, 2017). As societies became more sedentary and agricultural practices intensified, the process of urbanization began to emerge (Alday et al., 2018). Industrialization further accelerated urban growth, leading to the rise of cities and the need for cheap labour (Ortner & Frohlich, 2007). The transition from hunter-gatherer societies to agricultural settlements and then to industrial urbanization also had implications for the exposure of domesticated animals to urban environments, as they became integral to agricultural practices and urban life.

### significance of Transitions

The transition from foraging to farming and then to urbanization signifies radical changes in human diets, environments, and lifestyles, directly affecting the gut microbiome(Obregon-Tito et al., 2015)(Rosas-plaza et al., 2022). Agriculture introduced a narrower diet and close contact with domestic animals, changing microbial diversity and introducing new strains into the human gut. Urbanization further altered the microbiome with processed foods, antibiotics, and reduced microbial exposure, linking to the rise in modern diseases like obesity(Turnbaugh et al., 2008) and diabetes(Ussar et al., 2016). These shifts are crucial for understanding the microbiome’s role in health and disease.

### Comparative Perspective

Comparing these human transitions with animal domestication and urban adaptation provides insights into microbiome evolution. Domesticated and urban animals exhibit changes in their gut microbiota mirroring human developments, such as reduced diversity corresponding to consistent, less varied diets. Studying these parallels enhances our understanding of microbiota-host dynamics and the overarching impact of environment and diet on health across species(Jha et al., 2018).

#### Objectives

The primary aim of this paper is to elucidate the evolutionary trajectories of the gut microbiome in humans and animals as they transitioned from their ancestral, naturalistic environments to those heavily influenced by human activities. By examining the shift from foraging and wild habitats to agricultural practices and eventually to urbanized settings, this study seeks to unravel the complexities of microbiome diversity and composition changes. It will explore the convergent and divergent factors that drive these microbial shifts, offering a comparative analysis of the adaptive strategies within the gut ecosystems across species lines in response to human-induced environmental changes.

This paper posits that the analogous alterations in the gut microbiomes of humans and animals, as a consequence of transitioning from undisturbed to human-modified ecosystems, underscore a complex interaction of dietary, lifestyle, and environmental factors. These interactions are pivotal to decoding the evolution of host-microbe symbioses and understanding the broader implications for health and disease in the context of our rapidly changing world.

This paper is organized into several key sections to guide the reader through our comparative analysis systematically. We will begin with an exploration of historical transitions from hunter-gatherer lifestyles to agriculture in humans and the parallel process of animal domestication. the shared and unique aspects of microbiome evolution in humans and animals. The paper will conclude with a contemplation of the implications that these insights hold for future scientific inquiries, public health strategies, and policy-making considerations. Following this, we will delve into the methodologies employed to assess changes in the microbiome, providing a foundation for our results section. The discussion will integrate these findings, highlighting both.

## II. Methods

### A. Data Collection

The primary goal of the data collection process was to meticulously collect and organize alpha diversity indexes of both human and animal gut microbiomes from various ecological settings and evolutionary stages. This effort was critical for generating a robust dataset to allow for a comparative analysis of gut microbiome transformations as a result of urbanization and domestication transitions.

#### Quantitative Data Collection

##### Literature Review

A thorough review of existing literature was conducted in order to identify relevant research papers presenting data on gut microbe alpha diversity in humans and animals. Eleven research papers were carefully chosen for their relevance, methodological rigor, and clarity of the presented alpha diversity indexes. (Prabhu & Kamalakkannan, 2020)(Zhao et al., 2021)(Gibson et al., 2019)(Metcalf et al., 2017)(Sugden et al., 2020)(Maraci et al., 2022)(Colquhoun et al., 2019)(“Inside the Guts of the City: Urban-Induced Alterations of the Gut Microbiota in a Wild Passerine,” 2018)(Conteville et al., 2019)(Gomez et al., 2016)(Rosas-plaza et al., 2022)

##### Data Extraction

The alpha diversity indices reported in the selected papers were extracted manually and meticulously recorded in a structured Excel spread sheet. The extracted data included information about the alpha diversity indexes, the species and populations studied, and the ecological or evolutionary context in which the data was collected (for example, urban, domestic, wild, and so on).

##### Data Organization

To facilitate subsequent analyses, the Excel spread sheet was organized with separate sheets for human and animal data, as well as additional subdivisions based on ecological or evolutionary contexts. This well-organized dataset was used as the foundation for the quantitative analysis in this review.

#### □ Qualitative Data Collection

##### Literature Review

In addition to the papers chosen for quantitative data extraction, a large number of other research papers were reviewed to gather qualitative information about changes in gut microbiome compositions. The relevance of these papers to the themes of urbanization, domestication, and gut microbiome transformations was determined.

##### Data Extraction

From the selected papers, information about the microbial groups that increased or decreased, as well as other notable inferences about the implications of these changes, were extracted. Details on the functional roles of specific microbial groups, their relationships with host health, and the potential drivers of observed microbiome transformations were included.

##### Data Synthesis

The extracted qualitative data was synthesized in order to develop a comprehensive understanding of gut microbiome transformations and their implications. This synthesis aided the comparative and interpretative analyses conducted in this review, adding to the discussion of the parallel histories of human and animal gut microbiome evolutions during urbanization and domestication transitions.

### B. Quantitative Analysis

#### Exploratory Data Analysis (EDA)

The datasets were subjected to extensive exploratory data analysis with Python’s and Seaborn’s libraries. To visually represent the distribution of Shannon values across different environments and transitions, bar plots were used. The EDA’s primary goal was to not only understand the underlying patterns in the data but also to validate the effectiveness of the preprocessing steps and detect any potential outliers.

#### Statistical Analysis

Paired t-tests were used to determine the significance of observed changes in Shannon values, which were facilitated by the ‘stats’ module from Python’s ‘scipy’ package. This test was chosen because of its ability to compare two related samples, making it appropriate for our datasets in which Shannon values from one environment were compared to Shannon values from another for the same group of animals or areas. The p-values obtained from these t-tests were then used to infer the statistical significance of the observed changes. (Virtanen et al., 2020)

#### Correlation Analysis

To understand the relationships between initial gut microbiome diversity and the magnitude of change, a detailed correlation analysis was performed. Pearson’s correlation coefficient, calculated in Python with the ‘numpy’ library, assessed the strength and direction of the linear relationship between variables. This analysis revealed how the initial Shannon index in one environment could influence the magnitude of change when transitioning to another.(Harris et al., 2020)

#### Visualization

Visual aids were critical in interpreting and communicating the findings. Several plots, including bar plots with individual trend lines and box plots, were created in Python using the ‘matplotlib’ (Hunter, 2007) and ‘seaborn’ (Waskom, 2021) libraries. These visualizations helped to clarify data patterns and highlighted key transitions in gut microbiome diversity.

## III. Results

### A. Human Gut Microbiome Evolution

#### 1. Hunter-Gatherer Stage

Bacterial Composition: The gut microbiome of hunter-gatherer populations such as the Yanomami and BaAka is distinguished by a greater abundance of bacteria such as Prevotellaceae and Treponema (Conteville et al., 2019).

#### 2. Agricultural Stage

Bacterial Composition: As populations transitioned to agricultural lifestyles, the abundance of traditional microbiome features such as Prevotella and Treponema decreased noticeably. Instead, taxa belonging to the Firmicutes, such as Ruminococcaceae, Mogibacteriaceae, Faecalibacterium, and Leuconostoc, increased. (Rosas-plaza et al., 2022)(Gomez et al., 2016)

#### 3. Urban Stage

Bacterial Composition: The most significant changes in the gut microbiome were caused by urbanization. Bacteroidaceae increased significantly, which was linked to high fat consumption and the ability to degrade the mucus barrier in the gut. Bacteroidaceae and Rikenellaceae became more common, most likely as a result of their tolerance to bile, which has been linked to increased consumption of animal products(Filippo et al., 2010) (Obregon-Tito et al., 2015).

#### 4. Quantitative results

We discovered intriguing trends when we examined the Shannon index across three distinct societal groups: hunter-gatherers, agricultural, and urban. Hunter-gatherer communities had higher biodiversity indices, which increased as they transitioned to agricultural societies. However, this uptrend was not sustained when agricultural communities evolved into urban settings, revealing a decline in the Shannon index. The bar plot (Figure 1) visually encapsulated these patterns, highlighting the ebb and flow of biodiversity indices across societal transitions.

**Figure 1:**
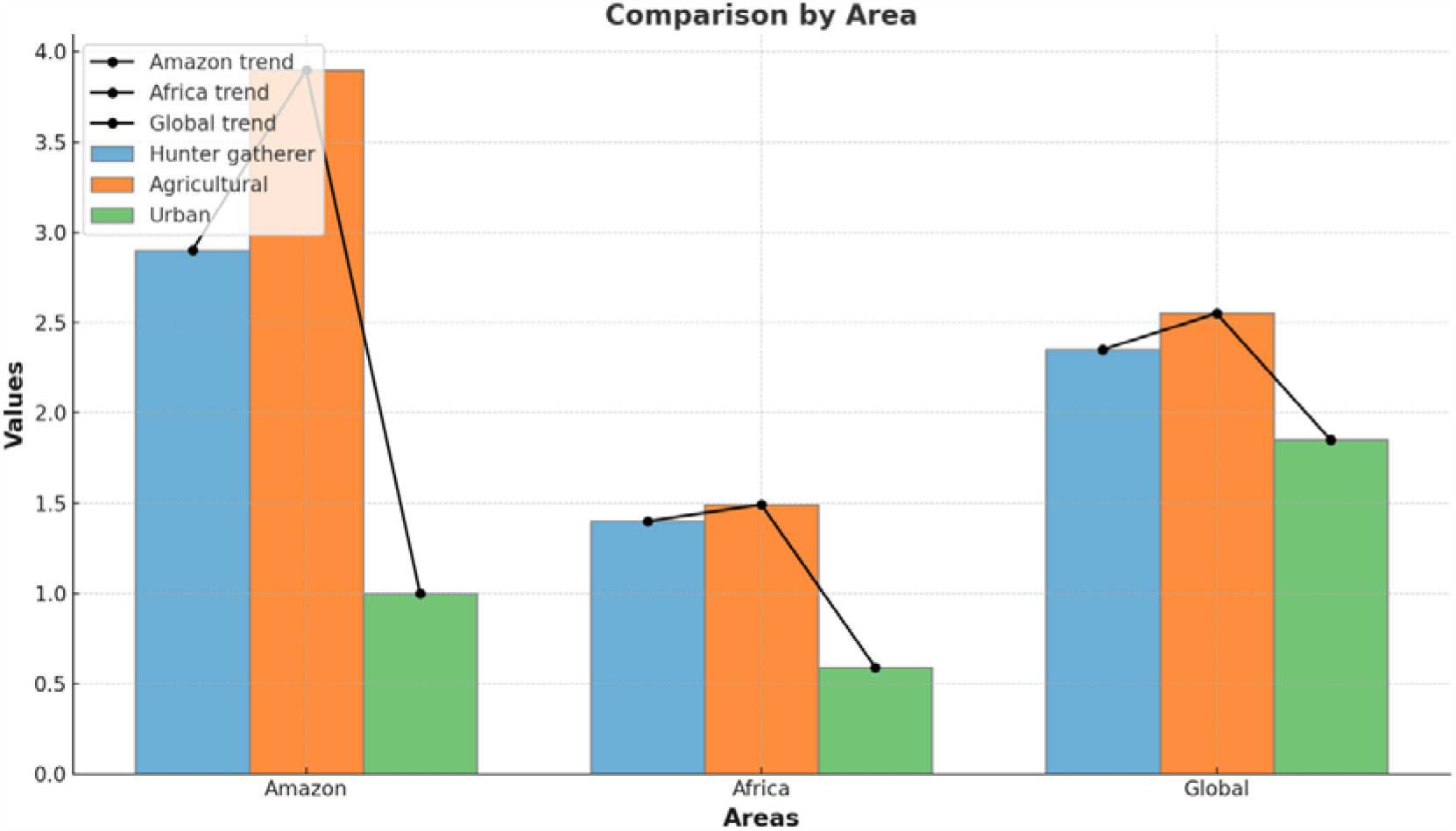
This graphical representation highlighted the complex relationship that exists between societal structures and ecological diversity.

**Figure 2:**
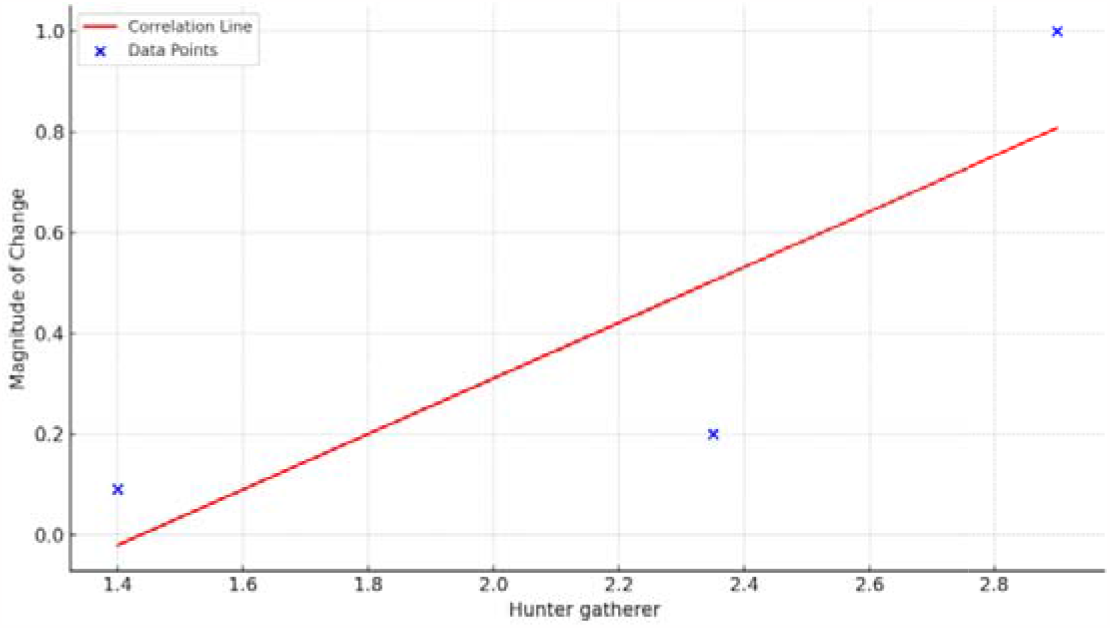
Correlation between”Initial Shannon index” and Magnitude of change in Shannon index Hunter-Gathering to Agricultural

In particular, we investigated the relationship between initial values and the magnitude of change in the context of the Shannon index.

##### Hunter-Gathering to Agricultural Transition

An initial examination of the data revealed a discernible positive trend across all areas between “Initial Shannon index” and Magnitude of change in Shannon index (). With a coefficient of r = 0.844, a strong positive correlation was discovered. This suggests that areas with higher Shannon indices for hunter-gatherers experienced a greater increase when agriculture was introduced.

##### Agricultural to Urban Transition

On the contrary, there was an inverse relationship between “Initial Shannon index” and Magnitude of change in Shannon index (Figure 3). Areas with higher Shannon indices in the agricultural phase, in particular, tended to have a smaller magnitude of change when transitioning to urban settings. This relationship’s correlation coefficient was (r = -0.859), indicating a strong negative trend.

**Figure 3:**
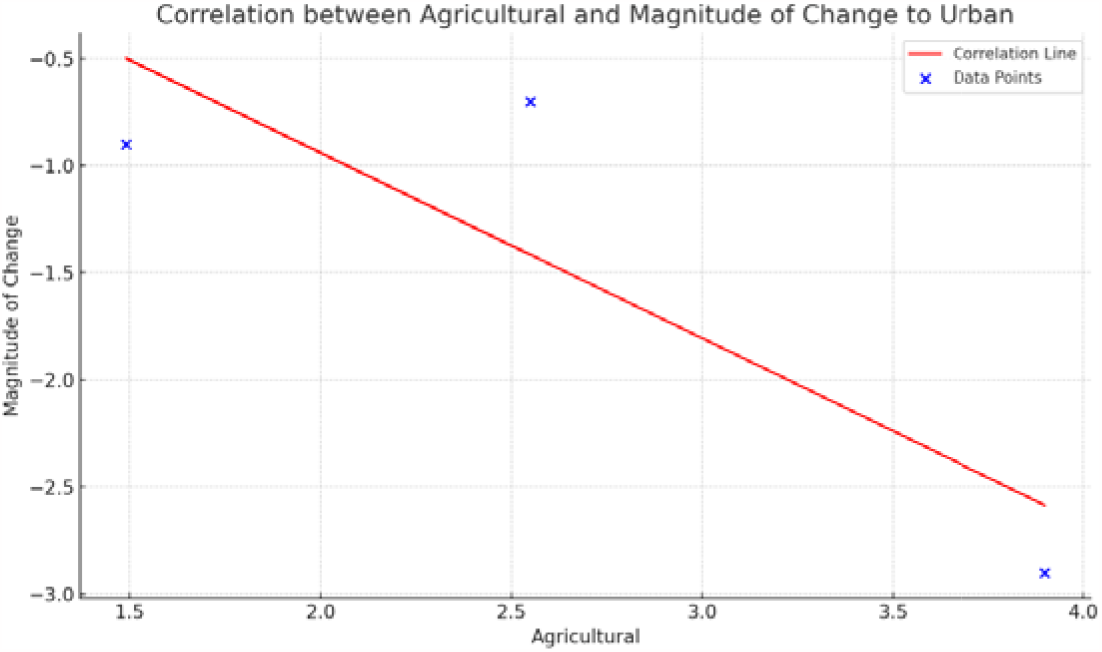

This disparity in trend between the two transitions highlights the complex interplay of factors influencing biodiversity and ecological structures as societies evolve and adapt.

### B. Animal Gut Microbiome Evolution

#### 1. Wild Stage

The initial wild stage of the gut microbiome, as observed in animals, represents a state of remarkable diversity and balance. This ecosystem, formed over millennia of co-evolution with nature, exemplifies a symbiotic relationship in which the host and its microbial inhabitants benefit from one another(Glazko et al., 2021). The Shannon values, which indicate microbiome diversity, are significantly higher in this wild setting, indicating a diverse range of microbial species. This diversity not only highlights the host’s adaptability to a variety of natural diets and environments, but it also highlights the gut ecosystem’s resilience to potential pathogens and external disruptions(Article, 2020).

#### 2. Domestication

Our research into the impact of domestication on gut microbiome diversity discovered significant differences in Shannon values between wild and domestic environments for the animals studied. The Shannon index was significantly higher in wild settings, averaging around 0.78, than in domestic settings, which had an average Shannon value of around 0.50. A paired t-test yielded a p-value of 0.0414, indicating that the decrease in gut microbial diversity caused by domestication was statistically significant.

A correlation analysis was performed to further elucidate the relationship between the initial microbial diversity in wild settings and the magnitude of its change upon domestication. The findings revealed a positive correlation of approximately 0.784, indicating that animals with a higher initial Shannon index in wild settings experienced a greater loss of diversity upon domestication. This suggests that domestication has a greater impact on the gut microbiome as microbial diversity increases (Figure 4).

**Figure 4:**
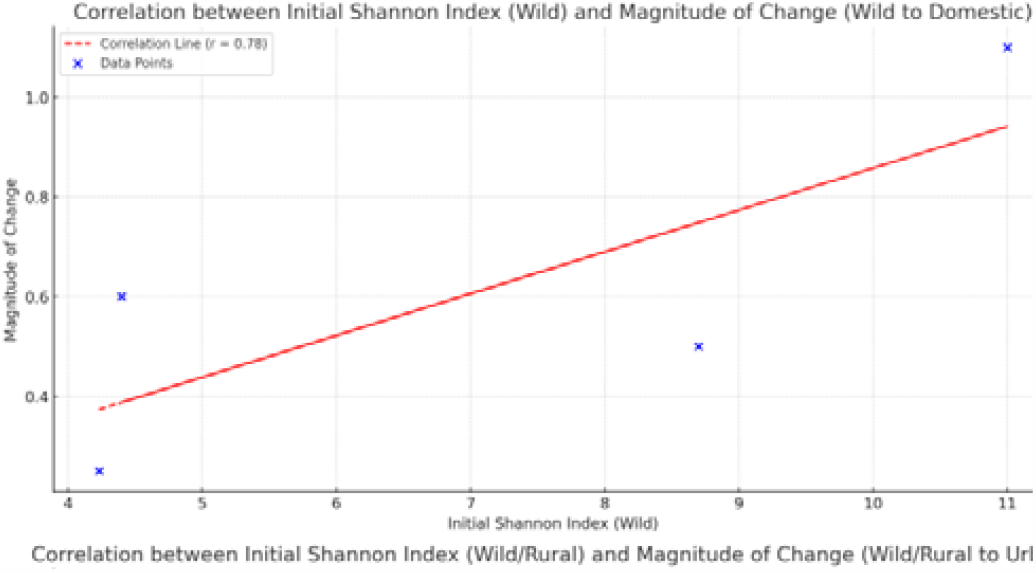
A correlation between the initial microbial diversity and the magnitude of its change due to domestication.

Our bar plots(Figure 5) with individual trend lines for each species emphasized these variations in the visual representation of this trend. Each trend line highlighted the shift in gut microbiome diversity as animals moved from wild to domestic environments, bolstering our numerical findings.

**Figure 5:**
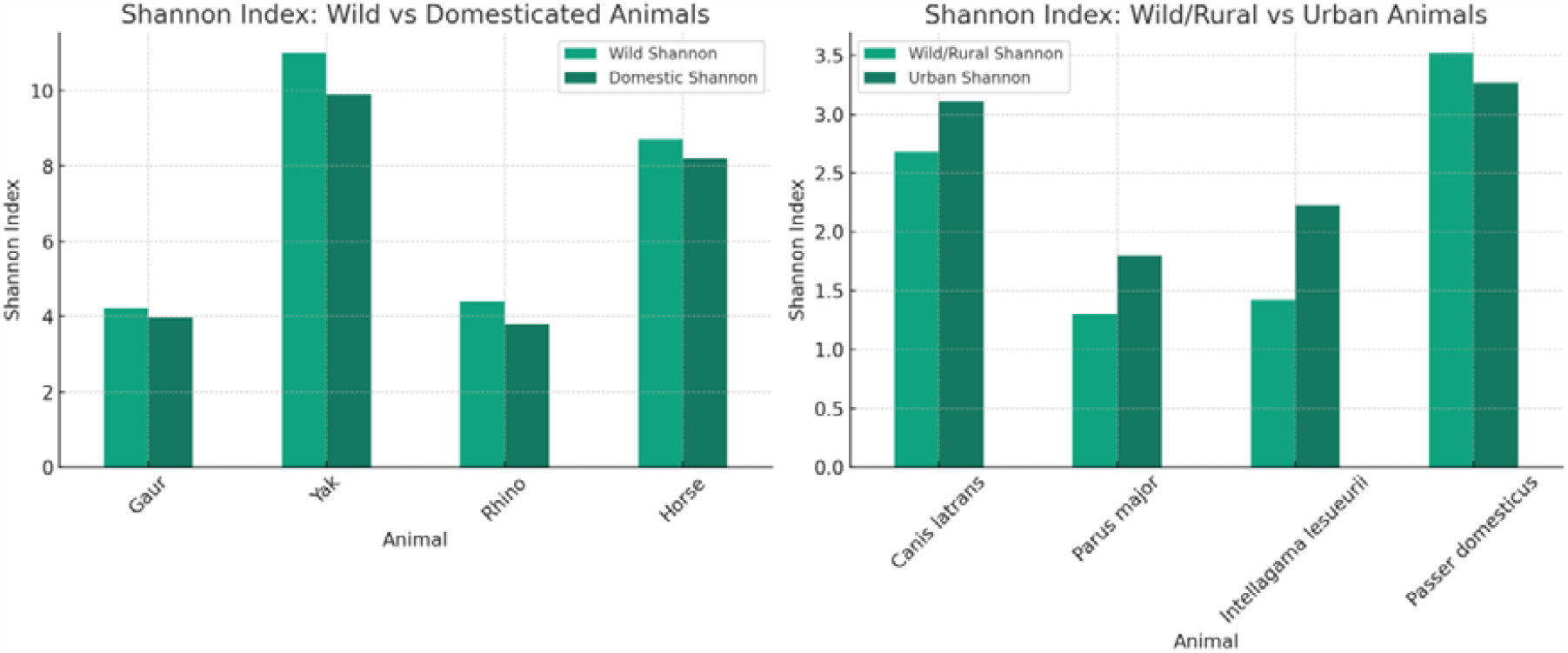
Bar plot comparing alpha diversity between Wild and Domestic animal (Left) and between Wild/Rural to Urban counterparts (Right)

## Compositional Chnage

### 1. Changes at the Phylum Level

#### Firmicutes

Firmicutes, a phylum of bacteria, has been observed to have a heightened abundance in domesticated animals. This observation has been made in various animal species, including mice (Ley et al., 2005), sifakas (Bensch et al., 2023), social subterranean rodents (Bensch et al., 2022), Tibetan sheep (Lv et al., 2021), ruminants (Faniyi et al., 2019), rabbits, and guinea pigs(Crowley et al., 2017).

#### Bacteroidetes

Bacteroidetes is a phylum of bacteria that has been found to decrease in domesticated animals. Several studies have reported a decrease in the abundance of Bacteroidetes in domesticated animals compared to their wild counterparts. For example, found that genetically obese mice (ob/ob) had a 50% reduction in the abundance of Bacteroidetes compared to lean mice (Ley et al., 2005). Similarly, observed an increased proportion of Firmicutes and a decrease in Bacteroidetes in obese mice (Schwiertz et al., 2010). In the context of domesticated animals, found a considerable decrease in the relative abundance of Bacteroidetes and a larger proportion of Firmicutes in obese animals compared to lean animals (Min et al., 2019). Additionally, found that the change from dominance by Actinobacteria in wild pigs to Bacteroidetes in domesticated pigs may be influenced by genetics, evolution, environmental change, and diet (Zhang et al., 2022).

#### Actinobacteria

Actinobacteria have been observed to increase in the gut microbiome during the domestication of animals. Several studies have investigated the effects of captivity and domestication on the gut microbiome and have found changes in the abundance of Actinobacteria. One study compared the gut microbiome of wild and captive mammals and found that the alpha diversity of gut bacteria remained consistent in some mammalian hosts, declined in others, and increased in captive rhinoceros (McKenzie et al., 2017). Another study examined the gut microbiota of wild and domesticated mammals, including chimpanzees and humans, and found a strong signal of domestication in the overall gut microbial community composition, with similar changes in composition observed with domestication and industrialization (Reese et al., 2021). Additionally, a study on the gut microbiome of neotropical bird species found that captivity status can have a significant effect on microbial composition, including the abundance of Actinobacteria (Hird et al., 2015).

#### Proteobacteria

Proteobacteria have been observed to increase in the gut microbiome during the domestication of animals. Comparative analyses of the gut microbiota of domesticated silkworms and their wild mulberry-feeding relatives revealed a highly diverse but distinctive gut microbiota in domesticated silkworms, dominated by the phyla Proteobacteria, Firmicutes, Actinobacteria, and Bacteroidetes (Chen et al., 2018). Additionally, a study comparing the gut microbiota of wild and domesticated mammals, including humans, found a strong signal of domestication in overall gut microbial community composition, with similar changes in composition observed during domestication and industrialization (Reese et al., 2021).

### 2. Modifications at the Class and Order Levels

#### Clostridia

Clostridia abundance has increased in domesticated animals, assisting in the fermentation of complex carbohydrates.(Pickard et al., 2017)

#### Bacteroidia

Depending on the diet, changes can be variable.(Turnbaugh et al., 2008)

#### Erysipelotrichi

Some increases in domesticated animals have been observed in specific studies.(Winter et al., 2013)

### 3. Urbanization

Our research into the effects of urbanization on gut microbiome diversity revealed significant differences between animals in wild, rural, and urban environments. Shannon values, which indicate microbiome diversity, were consistently higher in wild or rural settings, averaging around 0.67. In comparison, urban environments had a lower average Shannon value of around 0.42. This decrease in gut microbial diversity with urbanization was not only visible but also statistically significant, as evidenced by a p-value of 0.0077 in our paired t-test.

A correlation analysis was performed to investigate the relationship between initial microbial diversity in wild or rural settings and its change as animals transitioned to urban environments. The analysis produced a positive correlation of about 0.692. (Figure 6) This implies that animals with a higher initial Shannon index in wild or rural environments experienced a greater decline in diversity when they moved to cities.

**Figure 6:**
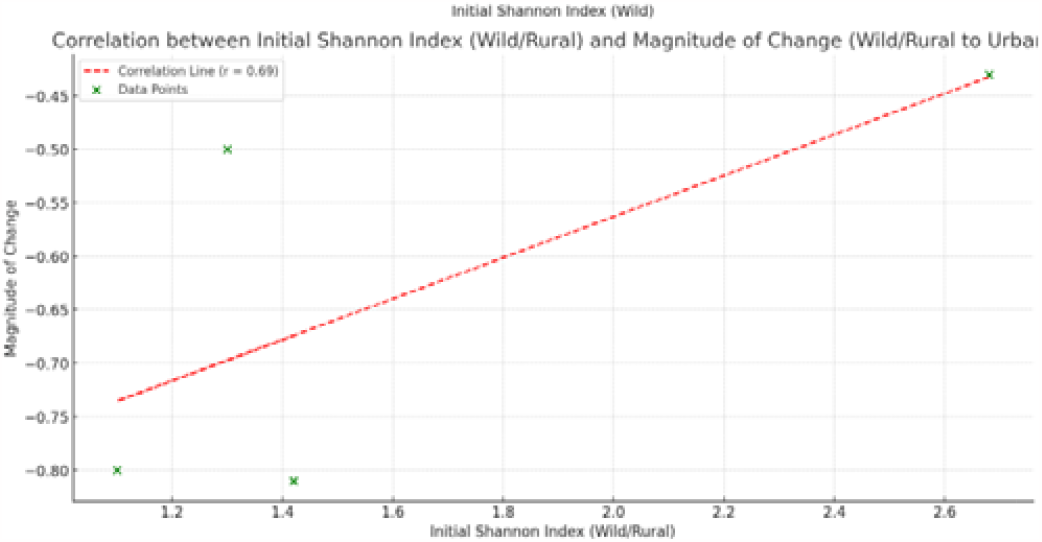
A correlation between the initial microbial diversity and the magnitude of its change due to Urbanization.

Graphical representations, specifically bar plots (Figure 5) with individual trend lines for each species, confirmed these findings visually. Each trend line clearly demonstrated the decrease in gut microbiome diversity as animals moved from wild or rural settings to urbanized environments, confirming our numerical findings.

#### Compositional Change

During urbanization, there are changes in the gut microbiome of animals compared to their wild counterparts. Several studies have investigated the alterations in specific microbial taxa during this process.

One important group of bacteria that has been studied is **Firmicutes**. David et al., 2013 found that the gut microbiome can be rapidly altered by diet, leading to changes in the abundance of Firmicutes. Additionally, Farinella et al., 2023 found that members of Firmicutes (specifically families Lactobacillaceae and Ruminococcaceae) decreased in relative abundance during peak infection of malaria in rhesus macaques.

Another group of bacteria that undergoes changes during urbanization is **Bacteroidetes**. Phoonlapdacha et al., 2022 reported that a specific type of resistant starch (RS4) can increase the abundance of Bacteroidetes in the gut microbiome. Conversely, Magne et al., 2020 suggests that a higher Firmicutes/Bacteroidetes ratio may be indicative of gut dysbiosis in obese patients.

Actinobacteria is another microbial group that can be affected by urbanization. Phoonlapdacha et al., 2022 found that RS4 significantly increased Actinobacteria in the gut microbiome. However, it is important to note that this study focused on pregnant women and the effects of specific types of foods.

Proteobacteria is a bacterial phylum that can also be influenced by urbanization. Farinella et al., 2023 observed a dramatic increase in Proteobacteria (specifically family Helicobacteraceae) during peak infection of malaria in rhesus macaques. This suggests that the presence of certain pathogens can lead to changes in the abundance of Proteobacteria.

In summary, during urbanization, the gut microbiome of animals can undergo changes in the abundance of specific microbial taxa. Firmicutes, Bacteroidetes, Actinobacteria, and Proteobacteria are among the bacterial groups that have been found to be affected. These changes can be influenced by factors such as diet, infection, and dysbiosis. Further research is needed to fully understand the mechanisms and implications of these alterations in the gut microbiome during urbanization.

## C. Comparative Analysis

### Hunter-Gatherer to Agricultural vs. Wild to Domesticated

In humans, the shift from hunter-gatherer to agricultural lifestyles led to a decrease in the diversity of the gut microbiome, particularly with a reduction in Prevotella and Treponema, and an increase in Ruminococcaceae and other Firmicutes.

Similarly, in the domestication of animals, there was a noted decrease in gut microbiome diversity, with an increase in Firmicutes and a decrease in Bacteroidetes. The underlying cause is analogous: a change from a varied, natural diet to a more consistent, restricted diet associated with agriculture or controlled feeding practices in domestication. In both scenarios, there is a transition from a diet and lifestyle that require a broader microbiome for the digestion of diverse fibrous foods, to one that can be managed by a less diverse microbial community, adapted to a narrower range of available nutrients.

### Agricultural to Urban vs. Rural to Urbanized Animals

For humans, urbanization saw a further shift with an increase in Bacteroidaceae, potentially due to dietary changes such as higher fat consumption, and a decline in overall microbial diversity.

In the case of urbanized animals, a similar decrease in diversity was noted, often with lower Shannon index values in urban settings compared to rural ones. Urbanization likely involves changes in diet due to processed foods, exposure to antibiotics, and less contact with natural environments, all factors that can reduce microbial diversity.

### Shared Factors

#### Diet

In both humans and animals, diet is a primary driver of gut microbiome composition. Changes from diverse, fibrous diets to more uniform, often processed diets are associated with reduced microbiome diversity in both groups.

#### Lifestyle

A sedentary lifestyle in urban settings compared to an active one in natural environments can influence the gut microbiome. Less physical activity can reduce gut motility and alter microbiota composition.

#### Antibiotic Exposure

Urbanization for both humans and animals often bring increased exposure to antibiotics, either through medical use or through food sources, which can significantly alter the gut microbiome.

### Divergent Factors

#### Genetic Adaptation

Humans have undergone specific genetic adaptations during their evolution that affect the gut microbiome differently from animals. For example, the ability to digest lactose into adulthood is a human-specific adaptation that can affect microbiome composition.

#### Environmental Control

Humans have more ability to control their environment compared to animals. This includes factors such as hygiene, climate control, and medical care, all of which can have distinct impacts on the gut microbiome.

#### Cultural Practices

Cultural elements, such as varied cooking methods, use of spices, and consumption of fermented foods, can diversify human microbiota in ways not paralleled in the animal kingdom.

#### Selective Breeding

Domesticated animals undergo selective breeding for certain traits, which can also select for certain microbiome profiles, a factor not at play in human populations.

## IV. Discussion

### Discussion

***Prevotella*** is a genus of bacteria that inhabits the human gut during the hunter-gatherer stage. Prevotella is known for its contribution in breaking down of carbohydrates and dietary fibers that are resistant to digestion by enzymes generally secreted in the human body (Precup & Vodnar, 2019). Prevotella secretes enzymes like cellulases and xylanases which has genetic potential to break down break down cellulose and xylan from foods (mostly plant-based) (Dubois et al., 2017). A few studies also pointed towards the potential role of Prevotella species as intestinal pathobionts [2]. Another microbe found in abundance in man during the hunter-gatherer phase was ***Treponoma***. Presence of Treponema in the gut microbiota of traditional rural populations indicates the presence of a bacterial community using <u>xylan, xylose</u> and <u>carboxymethylcellulose</u> to produce high levels of short-chain fatty acid (Flint et al., 2008). ***Firmicutes*** such as ***Ruminococcaceae, Mogibacteriaceae, Faecalibacterium, and Leuconostoc*** was increased in the agricultural phase of humans. These taxa of microbes too breakdown carbohydrates in the gut that can’t be digested by the body’s enzymes such as dietary fibre and resistant starch. The causes prebiotic fermentation on dietary fibres and produce metabolites like butyrate which further prevent inflammation of the intestinal lining. ***Firmicutes*** stimulates the production of hormones and natural oxidants like glutathione. The twain side of the presence of gut microbiota indicates towards obesity as found in many studies (Edermaniger, 2021). Ingestion of high-fibre foods in both the hunter-gatherer and agricultural phase resulted in presence of abundance of ***Firmicutes*** and ***Bacteroidetes***. In the urban set up, a rise in ***Rikenellaceae*** was observed which if present in abundance is associated with Lupus and Alzheimer’s disease in mice and colorectal cancer in humans (Wang et al., 2021)(Bello-Medina et al., 2022)(Hoang et al., 2022). An imbalance in these gut microbiota might even show implications Autoimmune Diseases, impaired barrier function and mucosal immune dysregulation (Wang et al., 2021).

***Firmicutes*** and ***Bacteroidetes*** are also found in abundance in the wild and domestic phase of animals’ gut respectively. The domestication of animals led to the presence of other gut microbes namely ***Actinobacteria and Proteobacteria***. The most lucrative prokaryotes in terms of both economics and biotechnology are actinobacteria, which are also the producers of bioactive secondary metabolites, including enzymes, antibiotics, antitumor agents, and immunosuppressive compounds (Citarasu, 2012). ***Actinobacteria*** also cause a range of other diseases in humans, including diphtheria (Corynebacterium diphtheria), Whipple’s disease (Tropheryma whipplei), and bacterial vaginosis (Gardnerella) (Lewin et al., 2016). ***Proteobacteria***, on the other hand, are believed to be essential for the gut’s preparation for colonisation by the stringent anaerobes needed for normal gut function because they consume oxygen and reduce the redox potential of the gut environment (Moon et al., 2018).

## VI. Conclusion

This investigation has revealed significant parallels in the shifts of gut microbiome composition between humans and animals as they transitioned from natural to anthropogenically influenced environments. Our findings underscore that both human and animal microbiomes have experienced marked alterations in diversity and function in response to dietary changes, sedentary lifestyles, and increased exposure to synthetic environments. In humans, the adoption of agriculture and later industrialization corresponded with reduced microbial diversity and the proliferation of microbiota associated with a range of chronic illnesses. Similarly, animals undergoing domestication and living in urban settings showed analogous patterns of microbial change, emphasizing the profound impact of human activity on evolutionary trajectories.

## VII. Data availability

All data tables of Shannon indexes of different animals and human indices in different geographical areas are available in this github repository with all the code for statistical analysis and plotting.

However, our study is not without limitations. The complexity of microbiome interactions, coupled with variations in genetic backgrounds and environmental exposures, means that our conclusions must be viewed as part of a larger mosaic of influencing factors. Furthermore, the retrospective analysis of historical transitions relies heavily on indirect evidence, such as archaeological findings and contemporary observations of remaining hunter-gatherer societies and wild animal populations.

For future research, there is a pressing need to develop more nuanced models that can account for the myriad of variables impacting microbiome evolution. Longitudinal studies tracking microbial dynamics over time in both humans and animals as they undergo environmental transitions would be particularly enlightening. Moreover, expanding our understanding of the gut-brain axis and its role in behavioral changes during these transitions could unlock new dimensions of the microbiome’s influence. Finally, as we look to mitigate the negative health impacts of these evolutionary changes, further exploration into the restoration of microbiome diversity through diet, probiotics, and policy-driven environmental changes will be crucial. Our study lays the groundwork for these endeavors, marking a path forward for those seeking to comprehend and harness the microbiome’s potential in shaping future health outcomes.

